# In Vivo mRNA Hacking with Staple Oligomers Prevents Myocardial Hypertrophy

**DOI:** 10.1101/2023.04.18.537290

**Authors:** Yousuke Katsuda, Takuto Kamura, Tomoki Kida, Takeru Saeki, Yua Itsuki, Yuri Kato, Taishi Nakamura, Motohiro Nishida, Yusuke Kitamura, Toshihiro Ihara, Masaki Hagihara, Shin-ichi Sato

## Abstract

The elucidation of gene-silencing mechanisms by RNA interference (RNAi) and antisense oligomers has drawn increasing attention to nucleic acid medicine. However, several challenges remain to be overcome, such as in vivo stability^1^, target selectivity ^2,3^, drug delivery^4,5^, and induced innate immunity^6^. Here, we report a new, versatile, and highly-selective method to hack RNA by controlling RNA structure using short oligonucleotides (RNA hacking: RNAh) in living cells. The oligonucleotide, named Staple oligomer, hybridizes specifically to a target mRNA and artificially induces an RNA higher-order structure, RNA G-quadruplex (RGq)^7^, on the mRNA. As a result, the RGq allows effective suppression of the target protein translation. This technology does not require cooperation with bioprocesses including enzymatic reactions as in RNAi or antisense technologies, permitting the introduction of artificial nucleic acids into Staple oligomers to increase their in vivo stability without compromising their effectiveness. The method was validated by translational regulation of the mRNAs of TPM3, MYD88, and TRPC6, in a cell-free system and in living mammalian cells. In vivo application of the technology to TRPC6 mRNA allowed us to prevent cardiac hypertrophy in transverse aortic constriction (TAC)-treated mice with no detectable off-target effects. This technology provides new insights into gene therapy after RNAi and antisense technologies.

## Main

Current developments in biotechnology have facilitated our understanding and elucidation of rare diseases whose treatment options are often limited. Meanwhile, developing small- and medium-sized molecular compounds that act as orphan drugs or therapeutic agents remains yet challenging. Today, gene therapy is a promising medical approach that allows for more precise and personalized treatment of serious and rare diseases since gene therapeutics can sequence-specifically target a disease-causing gene. Design of nucleic acid drugs as seed molecules can be easily and rationally done by referring to the sequence information of the target gene. Another advantage is that nucleic acid drugs can target DNA and RNA, unlike most conventional low-molecular-weight compounds. For example, the therapeutic RNA interference (RNAi) modality is used to suppress target gene expression in a sequence-specific manner through mRNA degradation or translational inhibition^8^. In 2008, Patisiran was approved as the first commercial RNA-interference-based therapeutics by the United States Food and Drug Administration and the European Commission for the treatment of hereditary amyloidogenic transthyretin amyloidosis with polyneuropathy in adults^9^. Nusinersen, an antisense oligonucleotide whose mechanism of action is exon inclusion, was recently developed and successfully demonstrated to have a substantial therapeutic effect on spinal muscular atrophy^10^, whose treatment has not been established yet.

Despite the aforementioned advantages, the development of nucleic acid drugs that suppress gene expression remains challenging. While siRNAs are the first choice in in-vitro studies for suppressing the expression of target genes, few nucleic acid drugs use the RNAi mechanism, partially due to their off-target effects and low stability in vivo^1-3,11^. Evidently, there is a huge gap between experimental and pharmaceutical technologies in use, making practical application difficult. Various attempts have been made to bridge the gap. For example, computational chemistry-based software is being developed that uses basic thermodynamic stability information to avoid off-target effects^12-16^. Meanwhile, since genetic information contains many sequences similar to the target gene, methods based on computational chemistry may be ineffective in the presence of homologous target genes. To prevent in vivo degradation, artificial nucleic acids with non-natural structures are introduced to nucleic acid drugs, such as BNA (bridged nucleic acid)^17,18^, GNA (glycol nucleic acid)^19-21^, TNA (threose nucleic acid)^22,23^, morpholino^24,25^, and other nucleic acid analogues with various substitutions in the ribose ring^26,27^. Although such non-natural nucleic acids remarkably stabilize nucleic acid drugs, nucleic acid-based therapeutics still pose a challenge because their intrinsic ability to suppress gene expression is greatly impaired by their incompatible structures for critical bioprocessing, including enzymatic reactions by Argonaute or RNaseH. More specifically, seed molecules for nucleic acid medicine can be developed in a short period of time, but the cost required for optimization is not much different from the cost for developing conventional small-molecule medicines.

Thus, developing a new technology that compensates for the shortcomings of conventional nucleic acid drugs would be a promising medical approach that allows more precise and personalized treatment of serious and rare diseases. Herein, we report a versatile and highly-selective technology to hack mRNA (RNA hacking: RNAh) using short nucleic acids, named Staple oligomers, by inducing a stable RNA G-quadruplex (RGq) structure that inhibits protein translation process.

### Mechanism of action of Staple oligomers in response to specific sequences of target RNAs

RGq is a thermodynamically stable nucleic acid structure that inhibits ribosome scanning in the 5’UTR of mRNA and represses protein translation. To form RGq, four consecutive guanine triplets (G_3_; G-tracts) are minimally required. Balasubramanian *et al*. reported that an RGq is formed in the genes whose sequences conform to the (G_3_-N_1-7_-G_3_-N_1-7_-G_3_-N_1-7_-G_3_) rule^28^. However, it was recently reported that genes with longer loop bases (N_7<_) also form RGq structures^29,30^. In this particular case, the loop region may form a higher-order structure, and the G-tracts are thought to be in spatial proximity to each other. This led to the idea that if a short nucleic acid that binds to the flanking sequences of two distant G-tracts is designed so as to draw the G-tracts in close proximity, specific RGq should be induced at the targeted locations in long RNA. Since the structures of RGq are very tight and ribosomes are stalled or arrested at RGq site in the 5’ UTR, Staple oligomers could work as a suppressor of the target gene (**Figure 1a**).

**Figure 1.**
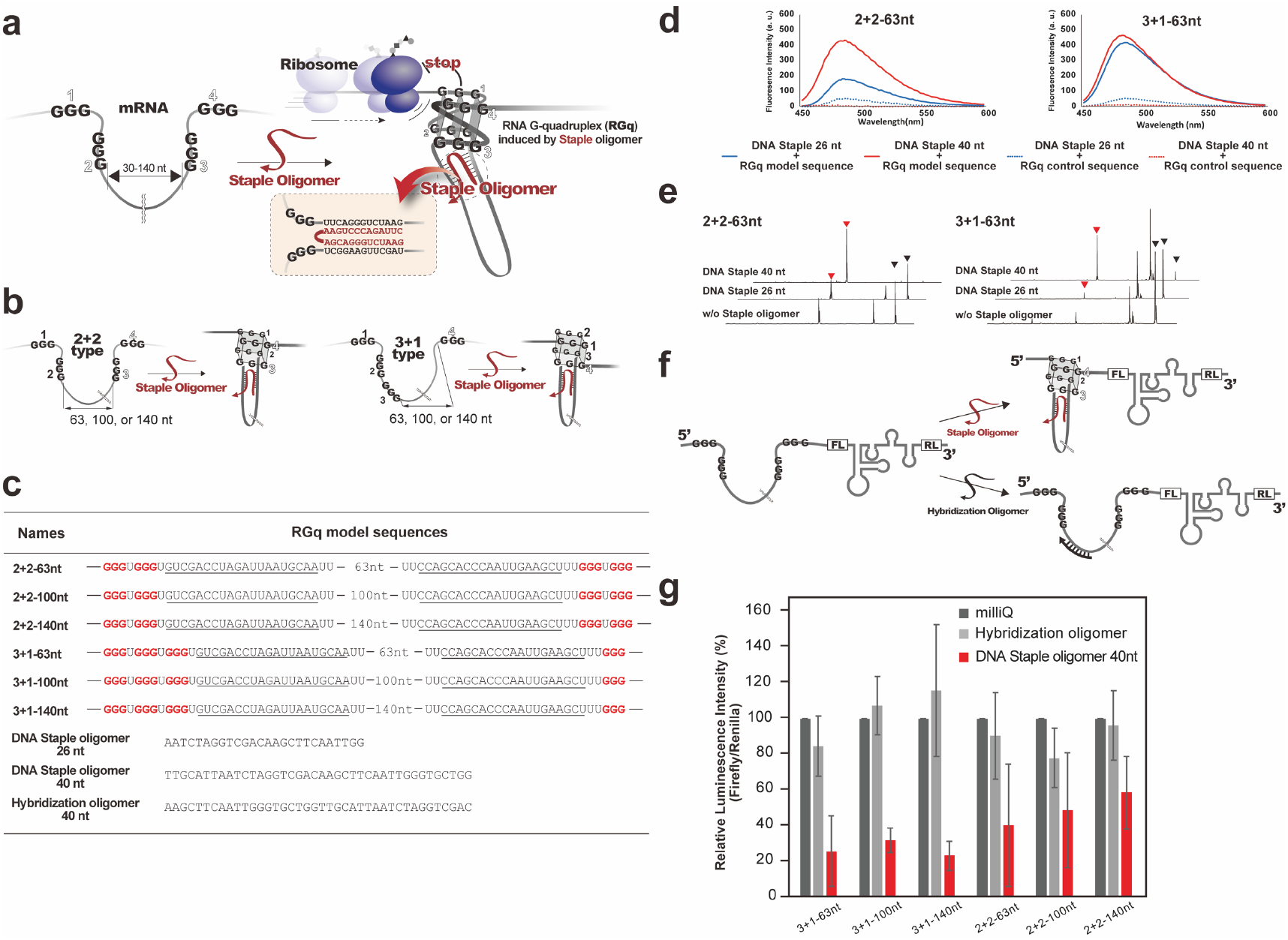
Characterizing RNAh technology. **a**. Mechanism of action of mRNA hacking. The technology requires two keys for unlocking to activate the mechanism: 1) strict recognition of two distinct target sequences with a Staple oligomer, and 2) induction of a stable RGq. The induced RGq inhibits ribosome scanning, resulting in translational suppression. **b**. RGq model sequences are illustrated. The RGq models have four G-tracts that are split into two each or one and three by 63-, 100-or 140-nt long loops. **c**. The sequences of RGq models, Staple oligomer, and Hybridization oligomer. The G-tracts are shown in red. Staple oligomer recognition sites are underlined. **d**. Evaluation of RGq formation with ThT. The blue and red curves show the fluorescence emission spectra of ThT in the presence of 26-nt or 40-nt DNA Staple oligomers, respectively. The solid and dashed curves represent the model sequence with G-tracts and the control with A-tracts, respectively. **e**. Identification of RGq formation on 2+2-63nt and 3+1-63nt RGq model sequences by Stop Assay. RTase-mediated cDNA synthesis was interrupted on all RGq model sequences in the presence of each DNA Staple oligomer. Red arrowhead indicates the arrest site of RTase with RGq induced by the Staple oligomers, and black arrowhead indicates RTase elongation end without RGq induction. **f**. Dual reporter genes encoding FL and RL in tandem were used to validate the mechanism of action of RNAh. RGq model sequence was placed in 5’UTR of FL. RL was placed downstream of an IRES as an internal standard. Staple oligomer induces RGq formation (upper panel), whereas Hybridization oligomer does not (lower panel). **g**. Effect of 40-nt DNA Staple oligomer on the translation of reporter RNA in vitro. The Staple oligomers significantly reduced translation from reporter RNA templates, whereas Hybridization oligomers did not.

One of the notable advantages of the new technology is that the addition of less toxic artificial nucleic acids to Staple oligomers does not degrade the drug’s efficacy because it needs no bioprocess to function. In addition, the present technology requires two keys for unlocking to activate the mechanism: 1) simultaneous hybridization of the Staple oligomers to the distant two target sequences, and 2) proximity of the split G-tracts to the target sequences, permitting to avoid off-target effects, a long-standing problem in the development of nucleic acid drugs. This high target specificity allows the technology to be applied to target highly homologous genes where computational chemistry cannot avoid off-target effects.

### Evaluation of RNAh technology

To evaluate RNAh technology in vitro, we prepared six model sequences with four G-tracts separated by a 63-, 100-, or 140-nt intervening loop sequence into 2+2 or 3+1 manner. In addition, two control sequences with A-tracts instead of G-tracts were also prepared (**Figure 1b, Figure 1c and Figure S1a**). First of all, RGq induction by Staple oligomers was verified by fluorimetry using the RGq probe thioflavin T (ThT), which shows enhanced emission around at 480-90 nm when bound to RGq^31^. To confirm the induction of RGq structure by Staple oligomers, we designed two DNA Staple oligomers with different lengths (26 or 40 nt). As a result, when the designed Staple oligomers were introduced into the RGq model sequences and the control sequences, remarkable fluorescent signals were successfully observed only in the presence of RGq model sequences, suggesting that RGq structure was induced by the Staple oligomers in all RGq model sequences (**Figure 1d and Figure S1b**).

RGq inhibits not only translation but also the elongation reaction of reverse transcriptase. Hagihara *et al*. developed the technique for RGq detection and location (StopAssay) that exploits this mechanism and reported various RGq identification experiments^32,33^. We further evaluated RGq induction activity of the Staple oligomers by the StopAssay. It showed that both of the Staple oligomers induced RGq formations at the expected position in the sequences of all RGq models (**Figure 1e**). This indicated that the RNAh technology with Staple oligomers can be universally used for various target RNAs.

We next evaluated the translation inhibition activity of Staple oligomers by in vitro translation. For the in vitro translation experiments, we prepared three dual reporter genes in which the translation of Firefly luciferase (FL) is controlled by the model RNAs with 63-, 100-, or 140-nt long loops at the 5’UTR, while Renilla luciferase (RL) translation mediated by internal ribosomal entry site (IRES) is used as internal standard (**Figure 1f**). Comparison of the luminescence signals from FL and RL allowed us to quantify the effects of the RGq structure in 5’UTRs on translation. The 40-nt DNA Staple oligomer significantly suppressed gene expression for all RGq model sequences, while the oligonucleotide that simply hybridized to the target (Hybridization oligomer) showed no detectable suppression of gene expression despite the length of it is the same with the Staple oligomers (**Figure 1g**). These results suggest that Staple oligomers would be a promising nucleic acid medicine that suppresses expression level of the target gene.

### Application of RNAh technology to TRPC6 Gene

Based on the evaluation of preliminary results, we applied the present technology to the myocardial hypertrophy-related TRPC6 (transient receptor potential cation channel subfamily C member 6) gene (**Figure 2a**)^34,35^, which is highly homologous to another gene of the same subfamily, TRPC3, and is difficult to target with existing nucleic acid-based technologies (**Figure S2**). For the initial trial, to confirm the induction of RGq by the Staple oligomer, we conducted a fluorescent study using ThT. When the Staple oligomer was added to the solution of 5’UTR of mouse TRPC6 mRNA, a remarkable fluorescent signal was observed, showing the formation of RGq (**Figure 2b**). The Staple-oligomer-length dependence of reverse transcription inhibition efficiency for the TRPC6 gene by StopAssay^32,33^. The results showed that the 26-nt long RNA Staple oligomer is sufficient for inhibition of reverse transcription (**Figure S3**).

**Figure 2.**
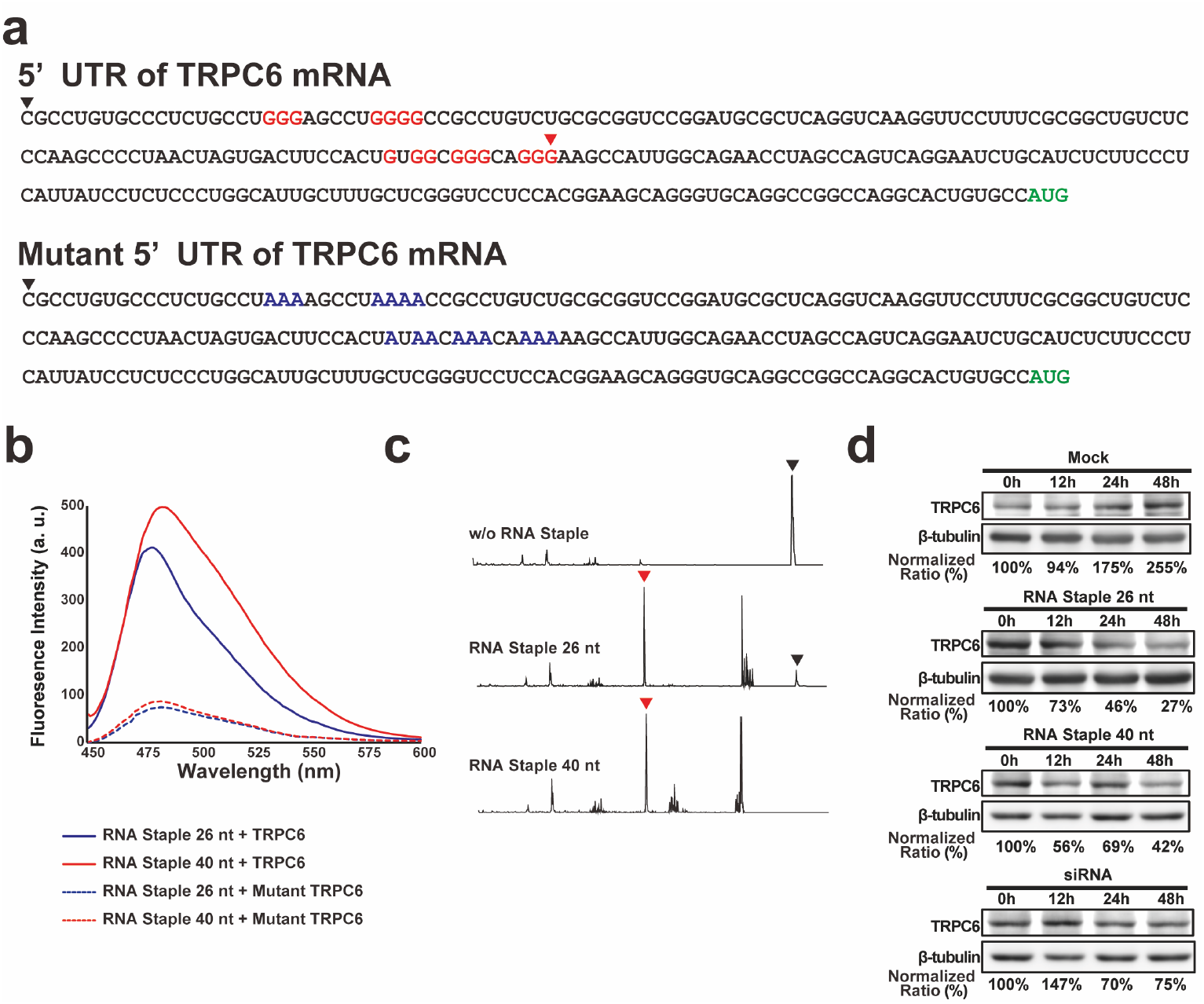
In vitro application of RNAh technology to the TRPC6 gene. **a**. Nucleotide sequences of the 5’UTR of TRPC6 mRNA and its synthetic control (mutant). The G-tracts are shown in red. The A-tracts, which replace the G-tracts, are shown in blue. Start codon (AUG) is shown in green. Red arrowhead indicates the arrest site of RTase with RGq induced by the Staple oligomer, and black arrowhead indicates RTase elongation end without RGq induction. **b**. Evaluation of RGq formation using ThT fluorescent probe. The red and blue curves show the fluorescence emission spectra of ThT in the presence of 26-nt or 40-nt RNA Staple oligomers, respectively. The solid and dashed curves represent the 5’UTR of TRPC6 and its mutant, respectively. **c**. Identification of RGq formation on the 5’UTR of TRPC6 mRNA by StopAssay. RTase-mediated cDNA synthesis was interrupted on the 5’UTR in the presence of each RNA Staple oligomer. Red and black arrowheads indicate the arrest sites of RTase with RGq induced by Staple oligomer and RTase elongation ends without RGq induction, respectively. **d**. Evaluation of the effects of RNA Staple oligomers and siRNA on TRPC6 expression in C2C12 cells by Western blotting. Protein expression levels were quantified relative to the expression of β-tubulin using ImageJ software.

The translation inhibition activity of the 26-nt and 40-nt RNA Staple oligomers was evaluated by in vitro translation. For the in vitro translation experiments, we prepared the two FL-RL dual reporter genes in which the translation of FL is controlled by the 5’UTR of TRPC6 mRNA (**Figure S4a**). Cell-free translation experiments of the reporter gene revealed that the Staple oligomers reduced the translational efficiency of the transcripts bearing the TRPC6 5’UTR by approximately 80%, while the Staple oligomers exhibited no detectable effects on the translation from its control sequence with A-tracts that is unable to form RGq (**Figure S4b**). These results are consistent with the results of fluorescent study using ThT for model sequences, showing that the Staple-oligomer-induced translational suppression is mediated by the induction of the RGq.

To validate the versatility of the RNAh technology, the possibility of regulating the expression of cancer-related genes, TPM3 and MYD88, was examined using the dual reporter genes incorporating their 5’ UTRs^36,37^. As a result, we succeeded in confirming the suppressive effect of RNAh on the expression of the cancer-related genes, demonstrating that RNAh is an extremely versatile technology (**Figure S4c, Figure S4d and Figure S4e**).

Based on the results, we applied the RNAh technology to endogenous TRPC6 mRNA in living C2C12 cells. The inhibition of TRPC6 expression in living cells by Staple oligomers was validated by Western blot analysis. For this experiment, short RNA expression vectors encoding 26-or 40-nt RNA Staple oligomers were employed to express within the cells. Western blotting revealed that even though TRPC6 expression tended to increase in C2C12 cells upon differentiation induction with horse serum (New Zealand origin), its expression decreased in a time-dependent manner in both cases where Staple oligomers were introduced (**Figure 2d and Figure S5a**). These results are consistent with those of the in vitro experiments.

The versatility of the RNAh technology in living cells was also demonstrated in its application to the cancer-related gene TPM3 (**Figure S5b**). The suppressive effect of the Staple oligomer was equal to or stronger than that of an siRNA which is known to be highly effective (**Figure 2d**)^38^.

### Application of RNAh technology in mice

We next applied the RNAh to the endogenous TRPC6 gene in mice. For this in vivo experiment, we constructed a system to express the 26-nt Staple oligomer in mice by introducing a short hairpin expression vector into adeno-associated virus 6 (AAV6) (**Figure 3a**). AAV6 is known to specifically infect the heart and liver and deliver the expression vector, but not the kidney^39,40^. AAV was administered to mice by intravenous tail injection. To evaluate the efficiency of expression repression by RNAh in each organ, the expression level of TRPC6 was verified by Western blot analysis. As expected, TRPC6 expression was significantly suppressed in the heart and liver, but no detectable change in TRPC6 expression was observed in the kidney (**Figure 3b**). Notably, no significant difference in the amount of TRPC6 mRNA was observed by quantitative PCR (qPCR) with or without the introduction of Staple oligomer by AAV6 (**Figure 3c**). A theoretical explanation for this observation is that siRNA functions by decreasing target mRNA, whereas the RNAh works by inducing a stable higher-order structure in the target mRNA without digesting the mRNA. The qPCR results provide strong evidence of the mechanism of action of the RNAh technology.

**Figure 3.**
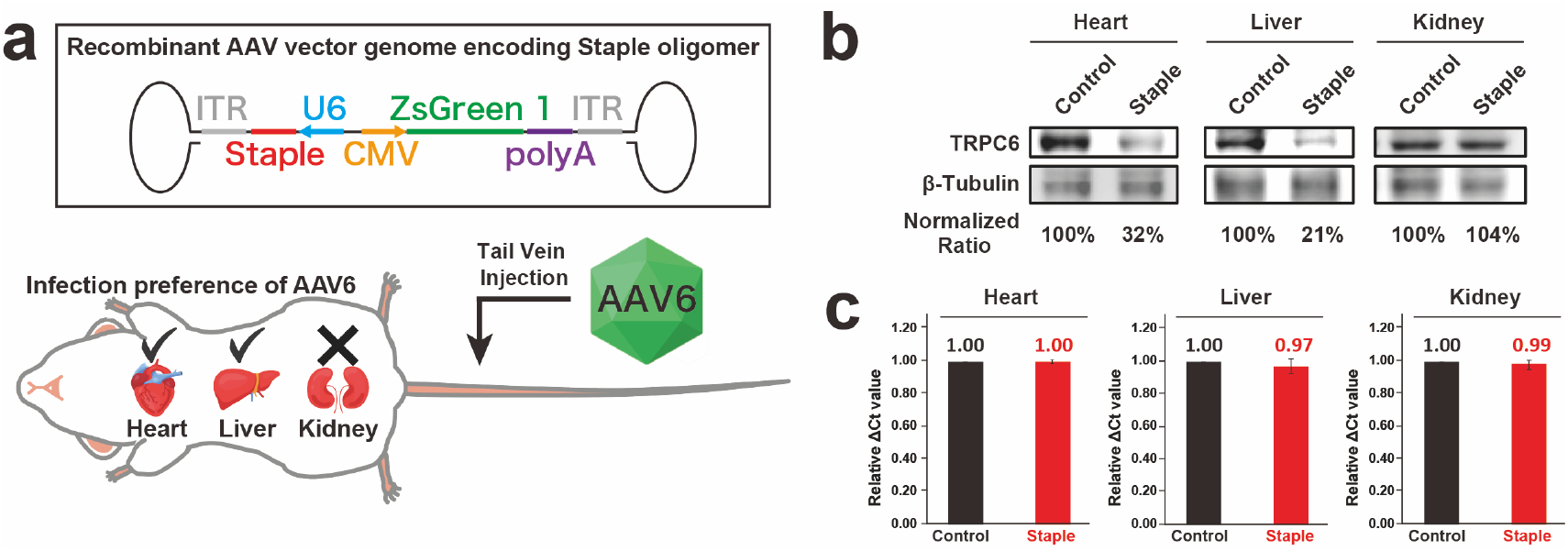
Effect of the Staple oligomers on TRPC6 gene expression in mice. **a**. Schematic of the recombinant AAV6 vector genome encoding the Staple oligomer. A short hairpin RNA expression vector was used to express the RNA Staple oligomer. The U6 promoter, Staple oligomer, and inverted terminal repeat (ITR) are shown in light blue, red, and gray, respectively. AAV6 infection is confirmed by ZsGreen (green) expression with CMV promotor (yellow). The RNA Staple oligomer is expressed by the AAV6 expression system in target organs. AAV6 infects the heart and liver, but not the kidney. **b**. Evaluation of protein expression levels of TRPC6 in each organ by Western blotting. TRPC6 expression was suppressed by 70-80% in the heart and liver by Staple oligomers, but not in the kidney, which is consistent with the infection-directedness of AAV6. **c**. Evaluation of TRPC6 mRNA expression levels in each organ by qPCR. No significant change in the mRNA expression levels was observed in each organ by introduction of RNA Staple oligomers, clearly supporting the action mechanism of RNAh that can suppress TRP6 expression without mRNA degradation.

### Prevention of myocardial hypertrophy in TAC-treated mice

TRPC6 gene is a potential therapeutic target for heart failure associated with myocardial hypertrophy^41-43^. To examine the therapeutic effects of Staple oligomers on cardiac hypertrophy, transverse aortic constriction (TAC)-treated mice, which overexpress TRPC6 and induce myocardial hypertrophy, were used as models (**Figure 4a**)^35,42^. The 26-nt Staple oligomers for RNAh were delivered to mouse hearts by AAV6 infection. As expected, the hearts from TAC-treated mice had significantly increased myocardial size and weight compared to those of mice without TAC treatment (sham). Remarkably, the hearts of TAC-treated mice treated with Staple oligomers, however, did not show as great of an increase in myocardial weight compared to those of mice in the group without Staple oligomer treatment (**Figure 4b and Figure S6a**). Echocardiographic evaluation of cardiac function showed no significant decrease in cardiac function after TAC treatment in the Staple oligomer-treated mice group, but a significant decrease in FS (fractional shortening) values, which measure the percentage change in left ventricular diameter during heart systole, in the non-Staple oligomer-treated mice (**Figure 4c and Figure S6b**). The results confirmed that Staple oligomers suppressed the decline in values reflecting cardiac function. We next evaluated the inhibitory effect of Staple oligomer on myocardial fibrosis by Masson’s trichrome staining. Staple oligomer had no effect on myocardial fibrosis in the sham group, whereas they significantly reduced the myocardial fibrosis in the TAC-treated mice (**Figure 4d**). Simultaneously, the expression level of TRPC6 was verified by Western blot analysis. TRPC6 expression was significantly suppressed by Staple oligomer in the hearts of TAC-treated mice (**Figure 4e**). We also evaluated the mRNA expression levels of cardiac hypertrophy marker genes ANP, BNP, and RcanI by qPCR in the TAC-treated hearts (**Figure 4f**). The expression of the selected markers was increased in TAC treatment, but Staple oligomer markedly suppressed their overexpression. These observations strongly suggest that Staple oligomer contribute to the maintenance of cardiac function in TAC-treated mice by suppressing TRPC6 expression.

**Figure 4.**
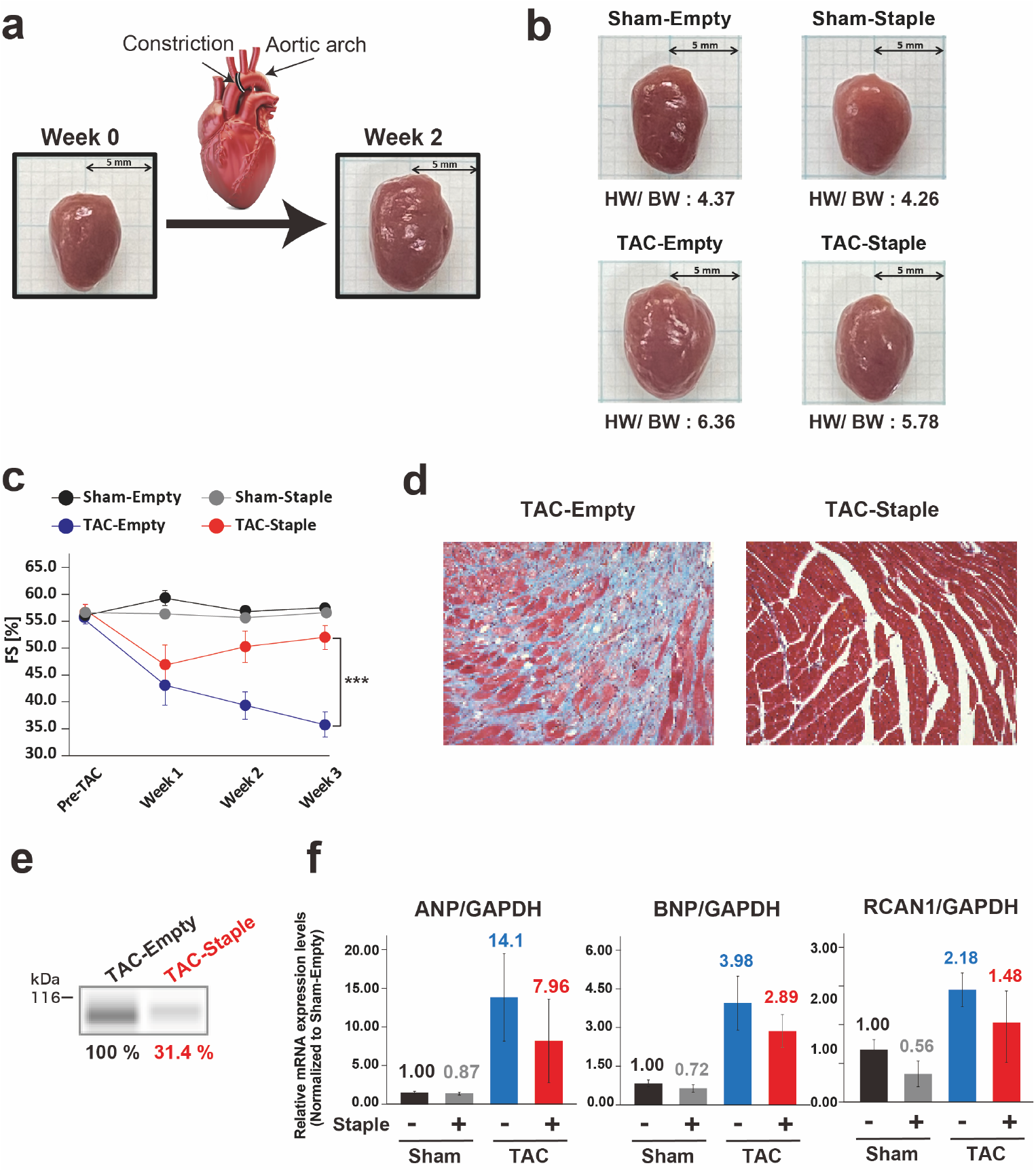
Effects of Staple oligomers on myocardial hypertrophy by transverse aortic constriction (TAC). **a**. Conceptual diagram of TAC. Stenosis of the aortic arch of the heart increases TRPC6 expression and causes myocardial hypertrophy. **b**. Representative pictures of hearts from Sham and TAC-treated mice infected with control-AAV6 or Staple-oligomer-AAV6. Myocardial enlargement of TAC-treated mice was significantly suppressed by the RNA Staple oligomer. **c**. Echocardiographic evaluation (FS values) revealed that cardiac function was maintained in the presence of the RNA Staple oligomers. Data are shown as mean ±SD. Statistical significance was determined by Student’s t-test: ^***^P<0.001, compared with control-AAV6 infection. **d**. Evaluation of myocardial fibrosis by Masson’s staining (blue area: area of fibrosis in the heart) showed that the Staple oligomer-treated myocardium was significantly less fibrotic. **e**. Evaluation of protein expression level of TRPC6 in mouse heart by Western blotting. TRPC6 expression was suppressed by approximately 70% with Staple oligomer treatment in the TAC-treated mouse hearts. **f**. Evaluation of mRNA expression levels of ANP, BNP, and RCAN1, known markers of cardiac hypertrophy, by qPCR. The mRNA expression levels were increased by TAC treatment, but the presence of Staple oligomers suppressed the increase in their mRNA expression levels.

### Microarray and proteomics analysis on myocardial hypertrophy

To evaluate the effects of Staple oligomers on gene expression, microarray analysis was performed in TAC-treated mice. In TRPC6, no detectable change was observed in the expression levels of its mRNA with the treatment of Staple oligomer. In contrast, the expression levels of cardiac fibrosis-related genes were significantly reduced with Staple oligomer treatments. These results strongly supported the suppressive effect on myocardial fibrosis indicated by Masson’s staining (**Figure S7a and Figure S7b**).

Proteomic analysis was conducted to prove the high target specificity of RNAh technology for target genes. The analysis revealed no detectable change in the expression levels of most genes, including computationally predicted homologous off-targets, such as Tns1 and Ehd2. Meanwhile, the expression levels of several genes were changed (**Figure S7c**). The changes may be experimental artifacts in proteomic analysis because of inconsistencies in the gene behaviours. For example, the gene Gsta4, which was identified as a significantly downregulated gene in the proteomic analysis, is known to affect the expression levels of the SOD2 and Gpx1 genes; however, these genes showed no significant change in their expression^44^. Thus, the proteomics observation led us to conclude that Staple oligomers suppress target gene expression with extremely high selectivity.

## Discussion

Here, we report RNAh technology as a promising and innovative nucleic acid-based therapeutics following RNAi and antisense technologies. Given the potential of RNAh technology, the future direction is to overcome issues inherent in nucleic acid medicine, such as in vivo stability, drug delivery, and off-target effects. Staple oligomer-mediated RNAh technology has a unique mechanism of action that is different from that of conventional nucleic acid drugs. Our technology does not require cooperation with bioprocesses such as enzymatic reactions, permitting the introduction of artificial nucleic acids into Staple oligomers to improve their stability in vivo without compromising their efficiency. Indeed, RGq was induced on model targets by DNA-, RNA-, and DNA/RNA-mixmer Staple oligomers, meaning that only the sequence-selective hybridization properties of Staple oligomers are crucial for Staple oligomers activity, but not the chemical properties of nucleotide units. The present study, however, leaves room for further discussion. The effective length of DNA Staple oligomers was different from that of RNA (**Figure S3**), possibly due to the differences in the intramolecular structure or duplex stability of each Staple oligomer. For drug discovery, optimization of base length would be a necessary process when installing artificial nucleotides in Staple oligomers.

The RNAh technology can be leveraged when a Staple oligomer accurately recognizes two sequences that apart from each other and subsequently induces an RGq structure on a target mRNA. RGq induction, however, requires a certain length of the Staple oligomer, implying that RGq formation should not occur even if a Staple oligomer binds to homologous off-targets with G-tracts. In length optimization of the Staple oligomer, its short hybridization significantly reduced the induction of RGq formation (**Figure S3**). Thus, this cooperative gene regulation technology combining target-specific hybridization and high-order structural induction, enables prescise, personalized, and safe gene therapy.

A common and critical challenge facing nucleic acid drugs, including RNAh technology, is drug delivery to target organs or cells. In the in vivo experiments reported here, RNA Staple oligomers highly expressed from infected AAV6 were used. The delivery of Staple oligomers to target organs and cells relies on the infection preference of AAV6; therefore, the delivery issue in the RNAh technology has not been solved in this stage. One possible solution to this issue is to combine the present technology with appropriate drug delivery systems^45-48^.

The new technology offers two advantages; (i) extremely high specificity should be expected because the RNAh requires both complementarity to a part of 5’UTR and nearby split G-tracts capable of forming RGq by proximity control, and (ii) artificial nucleic acids can be employed as Staple oligomers to increase in-vivo stability without diminishing their effectiveness. This technology, methodology-wise and therapeutics-wise, deserves the attention of nucleic acid-drug discovery with its high uniqueness, versatility, efficacy, and safety. Thus, this technology may make a meaningful impact on gene therapy for the precise and personalized treatment of serious and rare diseases.

## Supporting information

https://83.gigafile.nu/0727-c924b90087b1f4f28a461f33db56e8fa5

## ACKNOWLEDGMENT

This work was supported by JSPS (21J11197 to T. K., 22H02772 and 22K19395 to M. N., 20H02769 to T.I., 22K05326 to M. H., 20H02859, 21K18214 and 23H0282 to S.S., AMED under Grant Number JP20ak0101116, 23ak0101168, JP21lm0203004, JP22ym0126806, JP22ym0126813, JP20lm0203012 and 22fk0410055 (Y.K.), the Naito Foundation (T.I.), TERUMO Foundation for life Sciences and Arts (T.I.), ZE Research Program, IAE (ZE2022B-49 to Y.K. and ZE2022B-15 to S.S. and ZE2022-B22 to M.H.), ISHIZUE 2019 of Kyoto University Research Development Program (S.S.). This work was supported by JST FOREST Program, Grant Number JPMJFR211L.

