## Supplementary material for "In Vivo mRNA Hacking with Staple Oligomers Prevents Myocardial Hypertrophy": https://83.gigafile.nu/0727-c924b90087b1f4f28a461f33db56e8fa5

*[1.]Division of Materials Science and Chemistry, Faculty of Advanced Science and Technology, Kumamoto University, 2-39-1 Kurokami, Chuo-ku, Kumamoto 860-8555, Japan, [2.]StapleBio Inc., 2-39-1 Kurokami, Chuo-ku, Kumamoto 860-8555, Japan, [3.]Graduate School of Science and Technology, Hirosaki University, Hirosaki, Aomori 036-8561, Japan, [4.]Department of Physiology, Graduate School of Pharmaceutical Sciences, Kyushu University, Fukuoka 812-8582, Japan, [5.]Department of Medical Information Science, Graduate School of Medical Sciences, Kumamoto University, 1-1-1 Honjo, Chuo-ku, Kumamoto 860-8556, Japan, [6.]Department of Cardiovascular Medicine, Graduate School of Medical Sciences, Kumamoto University, 1-1-1 Honjo, Chuo-ku, Kumamoto 860-8556, Japan, [7.]Division of Cardiocirculatory Signaling, National Institute for Physiological Sciences & Exploratory Research Center on Life and Living Systems, National Institutes of Natural Sciences, Okazaki 444-8787, Japan, [8.]Institute for Chemical Research, Kyoto University, Uji, Kyoto 611-0011, Japan,*

*,*

### Supplementary Methods

#### Oligonucleotides

All oligonucleotides were purchased from Thermo Fisher Scientific or Eurofin Genomics. DNA oligonucleotides were used as Staple oligomers, primers and templates for PCR, and primers for qPCR. RNA oligonucleotides were used as Staple oligomers. Individual DNA and RNA oligonucleotides were stored at a concentration of 100 μM in TE buffer.

#### Thioflavin T (ThT) fluorescence measurement

A mixture of template RNA (4 μM), Staple oligomer (5 μM) in a buffer (20 mM Tris-HCl pH 7.8 and 100 mM KCl) was heated to 90 °C for 2 min, and then gradually cooled to 20 °C at a rate of 1.0 °C/min. For measurement, ThT (Sigma) was added to the mixture at 1 μM and incubated for 30 min in the dark at room temperature. Fluorescence intensity was measured with a spectrofluorometer FP-8500 (JASCO Corporation). The excitation wavelength was at 440 nm. The fluorescence spectra were obtained by taking the average of three scans made at 0.5 nm intervals from 450 to 600 nm. All of the fluorescence curves were normalized by subtraction of a control experiment without addition of a Staple oligomer.

#### StopAssay

A reaction mixture of template RNA (0.3 μM), Staple oligomer (1 μM), a 5'-FAM-labeled primer [5'-FAM- CGC CAG GGT TTT CCC AGT CAC GAC-3'] (0.1 μM) were heated to 90 °C for 2 min in a folding buffer (50 mM Tris-HCl pH 7.8, 150 mM KCl), and then cooled to 20 °C at a rate of 1.0 °C/min. ReverTra Ace reverse transcriptase (TOYOBO), $MgCl_2$ (5 mM), and dNTPs (0.83 mM) were then added to the reaction mixture, and the reaction was carried out at 42 °C for 30 min, and then the reaction was stopped by heating it at 99 °C for 5 min. After the reverse transcription, template RNAs were degraded by RNase H treatment. The produced cDNAs were analyzed on an SeqStudio Genetic Analyzer (Appliedbiosystems).

#### In vitro translation assay

A reaction mixture of template RNA (0.893 pmol), with or without Staple oligomer (1.116 pmol) and a buffer (50 mM Tris-HCl pH 7.8, 100 mM KCl) were heated to 90 °C for 2 min, gradually cooled to 20 °C at a rate of 1.0 °C/min. The samples were used as mRNA templates (0.893 pmol) in 20 μL of cell-free protein expression mixture (RTS 100

Wheat Germ CECF Kit, 5 Prime), with or without Staple oligomer (1.116 pmol), for 150 min at 24 °C. Luciferase activity was evaluated using the Luciferase Assay kit (Promega) and a POWERSCAN H1 microplate reader (BioTek).

**Cell cultures and Transfection**

C2C12 cells were maintained in medium A (Dulbecco's modified Eagle's medium, supplemented with 100-units/mL penicillin, 100-mg/mL streptomycin sulfate, and 10% (v/v) fetal bovine serum). MCF-7 cells were maintained in medium B (Eagle's minimum essential medium, supplemented with 100-units/mL penicillin, 100-mg/mL streptomycin sulfate, and 10% (v/v) fetal bovine serum). All cell lines were cultured at 37 °C in a humidified 5% $CO_2$ incubator. siRNA or Staple oligomer-expression vector was transfected into the cells using FuGENE HD (Promega) according to the manufacturer's instructions.

**Animal Studies**

Male BALB/c and C57BL/6J mice were purchased from Oriental Yeast Co., Ltd and CLEA JAPAN Inc, respectively. All mice were maintained under a 12:12-h light–dark cycle, and temperature were kept at 24 °C with ad libitum access to a regular chow diet and water. BLAB/c (6-week-old) or C57BL/6J (7-week-old) mice were used for in vivo translation assay or for phenotype evaluation, respectively. Pressure overload was produced by transverse aortic constriction (TAC). Sham-operated mice underwent the same operation, but without aortic constriction. All animal procedures were performed in accordance with Kumamoto University animal care guidelines and the Guide for the Care and Use of Laboratory Animals published by the by the U.S. National Institutes of Health (Publication No. 85-23, revised 1996).

**Adeno-associated virus (AAV) generation and Mouse transduction**

pAAV-U6-ZsGreen1 vector (5 μg) or Staple oligomer-expression construct (5 μg), pRC6 vector (5 μg) and pHelper vector (5 μg) were mixed for co-transfection of HEK293T cells in Optimem medium (Thermo Fisher Scientific) using TransIT-VirusGEN® Transfection Reagent (Mirus). 3 days after transfection, Staple-oligomer-expression or control AAVs were extracted from the cells and purified using AAVpro® Purification Kit Maxi (All Serotype) (Takara Bio) according to the manufacturer's protocol. The purified AAV titers were determined using AAVpro® Titration Kit Ver.2 (Takara Bio) for real time PCR according to the manufacturer's protocol. For tail vein injections of AAVs, the mice were anesthetized with isoflurane (pfizer). In vivo translation assay, BALB/c mice were injected in tail vain, with $1\times10^{9}$ vg of AAVs in 450 μL of PBS. In phenotypic evaluation, C57BL/6J mice were injected in tail vain, with $1.5\times10^{10}$ vg of AAVs in 450 μL of PBS.

**Echocardiography**

In vivo cardiac function was assessed by serial echocardiography in M-mode on Aplio300 (CANON MEDICAL SYSTEMS). Left ventricular internal dimension in diastole (LVDd), left ventricular internal dimension in systole (LVDs) and left ventricle fractional shortening were determined and then calculated by the average of two points from independently obtained M-mode images. All parameters were determined by an observer blinded to condition.

**Tissue collection and histology**

The heart and body weights of the sacrificed mice were recorded at the terminal point of the experiment. For pathological analyses, a mid-transverse cross-section of the heart encompassing both the left and right ventricle was dissected, fixed overnight in 4% paraformaldehyde, embedded in paraffin, cut into 3-μm-thick sections, and stained with Masson's Trichrome. The sections were analyzed using Axio Scope A1 microscope with Axiocam ERc 5s camera (ZEISS). Portions of the myocardium were frozen in liquid nitrogen and stored at -80 °C until used for Western blotting, RT-qPCR, proteomic and microarray analysis.

**Western blot analysis**

In cell translation assay, the cells were washed three times with cold PBS, and lysed with LIPA buffer (Nacalai Tesque) containing protease inhibitor cocktail (Nacalai Tesque). The cell lysates were passed 10 times through a 25G needle and centrifuged at 4 °C for 10 min. In vivo translation assay, the mouse heart, liver and kidney were homogenized with a lysis buffer (Cell Signaling Technology) containing protease inhibitor cocktail (Nacalai Tesque) by using μT-12 bead beater homogenizer (TITEC) and SLPe40 ultrasonic homogenizer (BRANSON). The lysates were centrifuged at 4 °C for 30 min to remove debris. The supernatants were transferred to new tubes and mixed with 6× sodium dodecyl sulfate (SDS) sample buffer (Nacalai Tesque), and then the mixture was heated at 95 °C for 5 min. The samples were separated on a 15% SDS-PAGE and blotted. The blotted protein bands were specifically visualized by antibodies against TRPC6 (16716: Cell Signaling Technology or 18236-1-AP: Proteintech) or β-Tubulin (66240-1-Ig: Proteintech), Signal Enhancer HIKARI for Western Blotting and ELISA (Nacalai

Tesque), and Chemi-Lumi One Super (Nacalai Tesque) on an ImageQuant LAS 500 (GE Healthcare).

**Western Blot Analysis using Abby Protein Simple System**

Simple western assay was performed in the Abby instrument (Protein Simple) using 12–230 kDa Separation 8×25 Capillary Cartridges (Protein Simple, SM-W004) according to the manufacturer's protocol. TRPC6 peak was detected by anti-TRPC6 antibody (Alomone Labs, ACC-017) used at 1:50 dilution and Anti-Rabbit Detection Module (Protein Simple, DM-001). A total protein assay using Total Protein Detection Module (Protein Simple, DM-TP01) and Replex Module (Protein Simple, RP-001) was also performed in the same run. TRPC6 peak area was determined using Compass software (Protein Simple) and normalized to that of total protein.

**RT-qPCR**

Total RNA was isolated from cells or mouse heart with ISOGEN (Nippon Gene) according to the manufacturer's protocol. First-strand cDNAs were prepared by reverse-transcription with an oligo (dT) primer (Thermo Fisher Scientific) and ReverTra Ace reverse transcriptase (TOYOBO) according to the manufacturer's protocol. The cDNAs were subjected to qPCR, using a pair of GAPDH primers to quantify mouse GAPDH mRNA [5'-AAC AGC AAC TCC CAC TCT TCC-3' and 5'-GTG GTC CAG GGT TTC TTA CTC -3'], a pair of TRPC6 primers to quantify mouse TRPC6 mRNA [5'-AAC TCG GGG AGA GAC TG-3' and 5'-ATA TGG CTT CAA GTG GAG-3'], a pair of ANP primers to quantify mouse ANP mRNA [5'-GGT CTA GTG GGG TCT TGC-3' and 5'-CGT CTG TCC TTG GTG CTG-3'], a pair of BNP primers to quantify mouse BNP mRNA [5'-CTG GGA CCA CCT TTG AAG TG-3' and 5'-GTG GCA AGT TTG TGC TCC-3'], a pair of RCAN1 primers to quantify mouse RCAN1 mRNA [5'- TCC AAA CCC TGT TTC CAG-3' and 5'-CTC TCC GGT TCA AAG TGC-3']. The qPCR analysis was performed on a MiniOpticon™ Real-Time PCR System (Bio-Rad Laboratories) with SYBR® Green (Takara Bio).

**Proteomic analysis by LC-MS/MS**

Total protein from the mouse heart was extracted in a 100-μL lysis buffer (pH 8.0) containing 10 mM Tris-HCl, 7 M urea, 2 M Thiourea, 5 mM magnesium acetate, 4%(w/v) of 3-[(3-Cholamidopropyl)-dimethylammonio]-1-propanesulfonate (CHAPS), and 1 tablet/50 mL of Complete™ Protease Inhibitor Cocktail Tablets. The extract mixture was disrupted using Sample Grinding Kit (Cytiva) with sonication, and subsequently concentrated using Amicon Ultra-0.5 and Ultracel-30 membranes (Millipore). The amounts of proteins in the concentrates were quantified by Bradford assay, and then 1.0 mg of total protein were fractionated according to the result of the assay. Thereafter, a 100 μL denaturing buffer containing 1.5 mg/mL of DTT and 100 mM of ammonium bicarbonate was added to the fractionated sample, and heated at 57 °C for 30 min. For protein alkylation, 100 μL of 10 mg/mL iodoacetamide (IAA) and 100 mM of ammonium bicarbonate was subsequently added to the mixture and incubated at room temperature for 30 min. Consecutively, 100 μL of Modified Trypsin (Promega) and 100 μL of 100 mM ammonium bicarbonate were added to the mixture, and incubated at room temperature for 30 min. Following the trypsin digestion, the resulting peptide sample was centrifuged at 20,000×g for 5 min to remove debris. Finally, 30 μL of 0.1% formic acid was added to the peptide sample, followed by vortexing and centrifugation at 20,000×g for 10 min. The supernatant was collected and used as a digested peptide sample for LC-MS/MS. LC-MS/MS was performed by Q Exactive™ Plus Hybrid Quadrupole-Orbitrap™ Mass Spectrometer (Thermo Scientific) coupled to UltiMate 3000 HPLC system (Thermo Scientific). Instrument operation and data acquisition were performed using Xcalibur Software (Thermo Scientific). The digested peptide samples were directly applied to the LC-MS/MS system. The samples were separated on a 500 × 0.075 mm capillary reversed-phase column (CERI) at a flow rate of 0.5 μL/min and a column temperature at 60 °C. Solvent A consisted of 0.1% formic acid (FA) in water, while solvent B consisted of 0.1% FA in acetonitrile (ACN). The following gradient was used for all samples:4% B for 0–5 min, 4–35% B from 5 to 340 min, 35–95% B from 340 to 350 min, 95% B until 360 min. The Q Exactive plus (Thermo Scientific) was operated in data dependent (dd) mode with full scans acquired at a resolution of 35,000 at 350 m/z and with dd-MS/MS scans acquired at a resolution of 17,500. The mass spectrometer was operated in positive mode in the scan range of 350-1500 m/z. Fixed first m/z is 150 in dd-MS/MS scans. Up to the top 10 most abundant isotope patterns with a charge ≥2 from the survey scan were selected with an isolation window of 1.6 m/z. The maximum ion injection times for the full scan and the dd-MS/MS scans were 60 ms and 100 ms respectively, and the automatic gain control (AGC) for the full scan and the dd-MS/MS scans were 3E6 and 1E5 respectively. Repeat sequencing of peptides was kept to a minimum by dynamic exclusion of the sequenced peptides for 20s.

**Data processing and analysis**

The database search was performed against the UniProt database of Mus musculus (Taxonomy ID:10090) using Proteome Discoverer ver. 2.3 (Termo Scientifc) and

MASCOT ver. 2.6 (Matrix Science) search engine software. Search parameters were as follows: static modifications, carbamidomethyl; dynamic modifications, oxidation (M); missed cleavages, up to 2; and MS/MS tolerance, 0.8 Da. The search result files were uploaded into Scaffold (version 4.11.1, Proteome Software) for protein identification and label-free quantitation based on an intensity-based absolute quantification (iBAQ) value. The two LC-MS/MS technical replicates (n = 2) of each of the two groups were combined in Scaffold.

**Microarray analysis**

Total RNA was extracted from mouse heart using ISOGEN (Nippon Gene) according to the manufacturer’s protocol and purified with After Tri Reagent RNA Clean Up Kit (Jena Bioscience Science). The total RNA (100 ng) and control RNA from One Color RNA Spike-In Kit (Agilent Technologies) were reverse transcribed according to the manufacturer’s recommendation. The resulting cDNAs were subsequently used for in vitro transcription and labeled with cyanine-3-labeled cytosine triphosphate using the Low RNA Input Amplification Kit (Agilent Technologies) according to the manufacturer’s protocol. Cy3-labeled cRNAs were then purified using the RNeasy Mini Kit (Qiagen) and checked for quality with the NanoDrop One (Thermo Fisher Scientific). 600 ng of Cy3-labeled cRNAs was hybridized to DNA microarray using SurePrint G3 Mouse GE Microarray 8 60K Ver. 2.0 (Agilent Technologies) at 65 °C and 10 rpm for 17 h. After hybridization, the arrays were washed with Gene Expression Wash Pack (Agilent Technologies), and then fluorescence from the array was scanned using an SureScan Microarray Scanner (Agilent Technologies). The accuracy of the relative quantification was validated using the One Color RNA Spike-In Kit. Normalization and data analyses were conducted using GeneSpring GX software ver. 14.9.1 (Agilent Technologies).

**Preparation of pUC19-TRPC6-5'UTR and pUC19-TRPC6-5'UTR-mutant constructs**

pUC19-TRPC6-5'UTR:

The mouse TRPC6-5'UTR DNA fragment was prepared by overlap extension PCR. Two double-stranded DNA (dsDNA) fragments (Section1 and Section2) were prepared by DNA polymerase reaction, using synthetic oligonucleotides [5'-TAA TAC GAC TCA CTA TAG GGC GCC TGT GCC CTC TGC CTG GGA GCC TGG GGC CGC CTG TCT GCG CGG TCC GGA TGC GC-3'] and [5'-CCA CAG TGG AAG TCA CTA GTT AGG GGC TTG GGA GAC AGC CGC GAA AGG AAC CTT GAC CTG AGC GCA TCC GGA CCG C-3'] (Section1), or synthetic oligonucleotides [5'-CTA GTG ACT TCC ACT GTG GCG GGC AGG GAA GCC ATT GGC AGA ACC TAG CCA GTC AGG AAT CTG CAT CTC TTC CC-3'] and [5'-GCA CCC CTG CTT CCG TGG AGG ACC CGA GCA AAG CAA TGC CAG GGA GAG GAT AAT GAG GGA AGA GAT GCA GAT TCC-3'] (Section2), respectively. Using the dsDNAs as templates, overlap extension PCR was performed to prepare the TRPC6-5'UTR DNA fragment, using a forward primer [5'-CGG TAC CCG GGG ATC TAA TAC GAC TCA CTA TAG GG-3'] and a reverse primer [5'-CGA CTC TAG AGG ATC GGC ACA GTG CCT GGC CGG CCT GCA CCC CTG C-3']. The PCR products were subcloned into the *Bam*H I site in pUC19 vector by using HiFi DNA Assembly Cloning Kit (New England Biolabs), then pUC19-TRPC6-5'UTR construct was obtained.

pUC19-TRPC6-5'UTR-mutant:

The mouse TRPC6-5'UTR-mutant DNA fragment was prepared by overlap extension PCR. Four dsDNA fragments (Section3, Section4, Section5 and Section6) were prepared by DNA polymerase reaction, using synthetic oligonucleotides [5'-TAA TAC GAC TCA CTA TAG GGC GCC TGT GCC CTC TGC CTA AAA GCC-3'] and [5'-GCG CAT CCG GAC CGC GCA GAC AGG CGG TTT TAG GCT TTT AGG CAG AG-3'] (Section3), [5'-GCG GTC CGG ATG CGC TCA GGT CAA GGT TCC TTT CGC GGC TGT CTC CC-3'] and [5'-GTT ATA GTG GAA GTC ACT AGT TAG GGG CTT GGG AGA CAG CCG CG-3'] (Section4), [5'-CTA GTG ACT TCC ACT ATA ACA AAC AAA AAA GCC ATT GGC AGA ACC TAG CC-3'] and [5'-GAG AGG ATA ATG AGG GAA GAG ATG CAG ATT CCT GAC TGG CTA GGT TCT GC-3'] (Section5), or [5'-CTC ATT ATC CTC TCC CTG GCA TTG CTT TGC TCG GGT CCT CCA CGG AAG C-3'] and [5'-GAG GAT CGG CAC AGT GCC TGG CCG GCC TGC ACC CCT GCT TCC GTG GAG G-3'] (Section6), respectively. Using each set of the dsDNAs Section3 and Section4, or Section5 and Section6, overlap extension PCR was performed, using a forward primer [5'-CGG TAC CCG GGG ATC TAA TAC GAC TCA CTA TAG

GG-3’] and a reverse primer [5’-GTT ATA GTG GAA GTC ACT AG-3’], or a forward primer [5’-CTA GTG ACT TCC ACT ATA AC-3’] and a reverse primer [5’-CGA CTC TAG AGG ATC GGC ACA GTG CCT GGC CGG CCT GCA CCC CTG C-3’], respectively. The two PCR products were used as templates for a subsequent overlap extension PCR. The overlap extension PCR was performed to prepare the TRPC6-5’UTR-mutant DNA fragment, using a forward primer [5’-CGG TAC CCG GGG ATC TAA TAC GAC TCA CTA TAG GG-3’] and a reverse primer [5’-CGA CTC TAG AGG ATC GGC ACA GTG CCT GGC CGG CCT GCA CCC CTG C-3’]. The PCR products were subcloned into the *Bam*H I site in pUC19 vector by using HiFi DNA Assembly Cloning Kit (New England Biolabs), and then pUC19-TRPC6-5’UTR-mutant construct was obtained.

**Preparation of pIRES-TRPC6-5’UTR-FL-RL and pIRES-TRPC6-5’UTR-mutant-FL-RL constructs**

pIRES-TRPC6-5’UTR-FL-RL:

The dsDNA for Renilla luciferase (RL) was amplified by PCR using psiCHECK-2 vector (Promega) as a template with a forward primer [5’-TCG ACC CGG GCG GCC ATG GCT TCC AAG GTG TAC G-3’] and a reverse primer [5’-TAA AGG GAA GCG GCC TTA CTG CTC GTT CTT CAG C-3’]. The PCR products were subcloned into the *Not* I site in pIRES vector (Clontech Laboratories) using the In-Fusion HD cloning kit, and then pIRES-RL construct was obtained. The dsDNA for firefly luciferase (FL) was amplified by PCR using psiCHECK-2 vector as a template with a forward primer [5’-CTA GCC TCG AGA ATT CGC CAC CAT GGC CGA TGC TAA GAA C-3’] and a reverse primer [5’-CTC GAC GCG TGA ATT TTA CAC GGC GAT CTT GCC-3’]. The PCR products were subcloned into the *Eco*R I site in pIRES-RL constructs using the In-Fusion HD cloning kit, and then pIRES-FL-RL construct was obtained.

pIRES-TRPC6-5’UTR-mutant-FL-RL:

The mouse TRPC6-5’UTR and TRPC6-5’UTR-mutant DNA fragments were prepared by overlap extension PCR. The TRPC6-5’UTR DNA fragment was amplified by PCR with a forward primer [5’-CGA CTC ACT ATA GGC TAG CCC GCC TGT GCC CTC TG-3’] and a reverse primer [5’-TCG GCC ATG GTG GCG AAT TCG GCA CAG TGC CTG G-3’] using pUC19-TRPC6-5’UTR construct as a template. The TRPC6-5’UTR-mutant DNA was amplified by PCR with a forward primer [5’-CGA CTC ACT ATA GGC TAG CCC GCC TGT GCC CTC TG-3’] and a reverse primer [5’-TCG GCC ATG GTG GCG AAT TCG GCA CAG TGC CTG G-3’] using pUC19-TRPC6-5’UTR-mutant construct as a template. Each PCR products was subcloned into the *Eco*R I site in pIRES-

FL-RL construct using HiFi DNA Assembly Cloning Kit (New England Biolabs), and then pIRES-TRPC6-5'UTR-FL-RL or pIRES-TRPC6-5'UTR-mutant-FL-RL constructs were obtained.

**Preparation of pMD19-RGq-model and pMD19-RGq-model-control constructs**

pMD19-RGq-model:

The RGq-model DNA fragment prepared by DNA polymerase reaction, using synthetic oligonucleotides [5'-TAA TAC GAC TCA CTA TAG ATT AGC ATA CGC TAC TGC AGA TGC GC-3'] and [5'-GCT GAA TAG CTT GCA AGT CAT GGC ATG CGC ATC TGC AGT AGC GT-3']. The dsDNA was subsequently amplified by PCR, using a forward primer [5'-TAA TAC GAC TCA CTA TAG-3'] and a reverse primer [5'-TCC AAC TAT GTA TAC CTG CTG AAT AGC TTG CAA GTC-3']. The PCR product was subcloned into the *Eco*R V site in pMD19 vector by using HiFi DNA Assembly Cloning Kit (New England Biolabs), then pMD19-RGq-model construct was obtained.

pMD19-RGq-model-2+2-63nt:

The 2+2-63nt DNA fragment was prepared by PCR using a synthetic oligonucleotide [5'-ACA CAG GAA ACA GCT ATG ACC ATG ATT ACG CCA AGT TTG CAC GCC TGC CGT TCG ACG ATT TAA TAC GAC TCA CTA TAG ATT AGC ATA CGC TAC TGC AGT GGG TGG GTG TCG ACC TAG ATT AAT GCA ATT CGT ACG AAG TTC ATA GCA TTT CCA GCA CCC AAT TGA AGC TTT GGG TGG GTG CAT GCC ATG ACT TGC AAG CTA TTC AGC AGG TAT ACA TAG TTG GAA ATC TCT GGA AGA TCC GCG CGT ACC GAG TTC TAA TTC ACT GGC CGT CGT TTT ACA ACG TCG TGA CTG GGA AAA CCC TGG CGT TAC CCA-3'] as a template with a forward primer [5'-TAA TAC GAC TCA CTA TAG A-3'] and a reverse primer [5'-TCC AAC TAT GTA TAC CTG-3']. As vector preparation, the pMD19-RGq-model construct was linearized by PCR using a forward primer [5'-CAG GTA TAC ATA GTT GGA AAT CTC TGG AAG ATC CG-3'] and a reverse primer [5'-CTA TAG TGA GTC GTA TTA AAT CGT CGA ACG GCA GG-3']. The 2+2-63nt DNA fragment was subcloned into the linearized pMD19-RGq-model vector using HiFi DNA Assembly Cloning Kit (New England Biolabs), and then pMD19-RGq-2+2-63nt construct was obtained.

pMD19-RGq-model-2+2-100nt:

The 2+2-100nt DNA fragment was prepared by PCR using pUC19 vector as a template with a forward primer [5'-CCT AGA TTA ATG CAA TTC GTG TGA AGA TCC TTT TTG ATA ATC TCA TGA C-3'] and a reverse primer [5'-TCA ATT GGG TGC TGG AAA TGA ACT CAC GTT AAG GGA TTT TGG TCA TGA G-3']. As vector preparation, the pMD19-RGq-model-2+2-63nt construct was linearized by PCR using a forward primer [5’-CAT TTC CAG CAC CCA ATT GAA GCT T-3’] and a reverse primer [5’-ACG AAT TGC ATT AAT CTA GGT CGA C-3’]. The 2+2-100nt DNA fragment was subcloned into the linearized pMD19-RGq-model-2+2-63nt vector using HiFi DNA Assembly Cloning Kit (New England Biolabs), and then pMD19-model-2+2-100nt construct was obtained.

pMD19-RGq-model-2+2-140nt:

The 2+2-140nt DNA fragment was prepared by PCR using pUC19 vector as a template with a forward primer [5’-CCT AGA TTA ATG CAA TTC GTG TGA AGA TCC TTT TTG ATA ATC TCA TGA C-3’] and a reverse primer [5’-TCA ATT GGG TGC TGG AAA TGT TTG ATC TTT TCT ACG GGG TCT GAC GCT CAG-3’]. The 2+2-140nt DNA fragment was subcloned into the linearized pMD19-RGq-model-2+2-63nt vector using HiFi DNA Assembly Cloning Kit (New England Biolabs), and then pMD19-RGq-model-2+2-140nt construct was obtained.

pMD19-RGq-model-3+1-63nt:

The 3+1-63nt DNA fragment was prepared by PCR using a synthetic oligonucleotide [5’-ACA CAG GAA ACA GCT ATG ACC ATG ATT ACG CCA AGT TTG CAC GCC TGC CGT TCG ACG ATT TAA TAC GAC TCA CTA TAG ATT AGC ATA CGC TAC TGC AGT GGG TGG GTG GGT GTC GAC CTA GAT TAA TGC AAT TCG TAC GAA GTT CAT AGC ATT TCC AGC ACC CAA TTG AAG CTT TGG GTG CAT GCC ATG ACT TGC AAG CTA TTC AGC AGG TAT ACA TAG TTG GAA ATC TCT GGA AGA TCC GCG CGT ACC GAG TTC TAA TTC ACT GGC CGT CGT TTT ACA ACG TCG TGA CTG GGA AAA CCC TGG CGT TAC CCA-3’] as a template with a forward primer [5’-TAA TAC GAC TCA CTA TAG AA-3’] and a reverse primer [5’-TCC AAC TAT GTA TAC CTG-3’]. The 3+1-63nt DNA fragment was subcloned into the linearized pMD19-RGq-model vector using HiFi DNA Assembly Cloning Kit (New England Biolabs), and then pMD19-RGq-3+1-63nt construct was obtained.

pMD19-RGq-model-3+1-100nt:

The 3+1-100nt DNA fragment was prepared by PCR using pUC19 vector as a template with a forward primer [5’-CCT AGA TTA ATG CAA TTC GTG TGA AGA TCC TTT TTG ATA ATC TCA TGA C-3’] and a reverse primer [5’-TCA ATT GGG TGC TGG AAA TGA ACT CAC GTT AAG GGA TTT TGG TCA TGA G-3’]. As vector preparation, the pMD19-RGq-model-3+1-63nt construct was linearized by PCR using a forward primer [5’-CAT TTC CAG CAC CCA ATT GAA GCT T-3’] and a reverse primer [5’-ACG AAT TGC ATT AAT CTA GGT CGA C-3’]. The 3+1-100nt DNA fragment was subcloned into the linearized pMD19-RGq-model-3+1-63nt vector using

HiFi DNA Assembly Cloning Kit (New England Biolabs), and then pMD19-model-3+1-100nt construct was obtained.

pMD19-RGq-model-3+1-140nt:

The 3+1-140nt DNA fragment was prepared by PCR using pUC19 vector as a template with a forward primer [5'-CCT AGA TTA ATG CAA TTC GTG TGA AGA TCC TTT TTG ATA ATC TCA TGA C-3'] and a reverse primer [5'-TCA ATT GGG TGC TGG AAA TGT TTG ATC TTT TCT ACG GGG TCT GAC GCT CAG-3']. The 3+1-140nt DNA fragment was subcloned into the linearized pMD19-RGq-model-3+1-63nt vector using HiFi DNA Assembly Cloning Kit (New England Biolabs), and then pMD19-RGq-model-3+1-140nt construct was obtained.

pMD19-RGq-model-control-2+2-63nt:

The 2+2-63nt-control insert DNA was prepared by PCR using a synthetic oligonucleotide [5'-ACA CAG GAA ACA GCT ATG ACC ATG ATT ACG CCA AGT TTG CAC GCC TGC CGT TCG ACG ATT TAA TAC GAC TCA CTA TAG ATT AGC ATA CGC TAC TGC AGT AAA TAA ATG TCG ACC TAG ATT AAT GCA ATT CGT ACG AAG TTC ATA GCA TTT CCA GCA CCC AAT TGA AGC TTT AAA TAA ATG CAT GCC ATG ACT TGC AAG CTA TTC AGC AGG TAT ACA TAG TTG GAA ATC TCT GGA AGA TCC GCG CGT ACC GAG TTC TAA TTC ACT GGC CGT CGT TTT ACA ACG TCG TGA CTG GGA AAA CCC TGG CGT TAC CCA-3'] as a template with a forward primer [5'-TAA TAC GAC TCA CTA TAG AA-3'] and a reverse primer [5'-TCC AAC TAT GTA TAC CTG-3']. As a vector preparation, the pMD19-RGq-model construct was linearized by PCR, using a forward primer [5'-CAG GTA TAC ATA GTT GGA AAT CTC TGG AAG ATC CG-3'] and a reverse primer [5'-CTA TAG TGA GTC GTA TTA AAT CGT CGA ACG GCA GG-3']. The 2+2-63-nt-control DNA fragment was subcloned into the linearized pMD19-RGq-model vector using HiFi DNA Assembly Cloning Kit (New England Biolabs), and then pMD19-model-2+2-63nt control construct was obtained.

pMD19-RGq-model-control-2+2-100nt:

The 2+2-100nt-control insert DNA was prepared by PCR using pUC19 vector as a template with a forward primer [5'-CCT AGA TTA ATG CAA TTC GTG TGA AGA TCC TTT TTG ATA ATC TCA TGA C-3'] and a reverse primer [5'-TCA ATT GGG TGC TGG AAA TGA ACT CAC GTT AAG GGA TTT TGG TCA TGA G-3']. As a vector preparation, pMD19-2+2-63nt-control construct was linearized by PCR, using a forward primer [5'-CAT TTC CAG CAC CCA ATT GAA GCT T-3'] and a reverse primer [5'-ACG AAT TGC ATT AAT CTA GGT CGA C-3']. The 2+2-100-nt-control DNA fragment was subcloned into the linearized pMD19-2+2-63nt-control vector using HiFi DNA Assembly Cloning Kit (New England Biolabs), and then pMD19-model-2+2-100-nt-control constructs were obtained.

pMD19-RGq-model-control-2+2-140nt:
The 2+2-140nt-control insert DNA was prepared by PCR using pUC19 vector as a template with a forward primer [5'-CCT AGA TTA ATG CAA TTC GTG TGA AGA TCC TTT TTG ATA ATC TCA TGA C-3'] and a reverse primer [5'-TCA ATT GGG TGC TGG AAA TGT TTG ATC TTT TCT ACG GGG TCT GAC GCT CAG-3']. The 2+2-140-nt-control DNA fragment was subcloned into the linearized pMD19-2+2-63nt-control vector using HiFi DNA Assembly Cloning Kit (New England Biolabs), and then pMD19-model-2+2-140-nt-control constructs were obtained.

pMD19-RGq-model-control-3+1-63nt:
The 3+1-63nt-control insert DNA was prepared by PCR using a synthetic oligonucleotide [5'-ACA CAG GAA ACA GCT ATG ACC ATG ATT ACG CCA AGT TTG CAC GCC TGC CGT TCG ACG ATT TAA TAC GAC TCA CTA TAG ATT AGC ATA CGC TAC TGC AGT AAA TAA ATA AAT GTC GAC CTA GAT TAA TGC AAT TCG TAC GAA GTT CAT AGC ATT TCC AGC ACC CAA TTG AAG CTT TAA ATG CAT GCC ATG ACT TGC AAG CTA TTC AGC AGG TAT ACA TAG TTG GAA ATC TCT GGA AGA TCC GCG CGT ACC GAG TTC TAA TTC ACT GGC CGT CGT TTT ACA ACG TCG TGA CTG GGA AAA CCC TGG CGT TAC CCA-3'] as a template with a forward primer [5'-TAA TAC GAC TCA CTA TAG AA-3'] and a reverse primer [5'-TCC AAC TAT GTA TAC CTG-3']. As a vector preparation, the pMD19-RGq-model construct was linearized by PCR, using a forward primer [5'-CAG GTA TAC ATA GTT GGA AAT CTC TGG AAG ATC CG-3'] and a reverse primer [5'-CTA TAG TGA GTC GTA TTA AAT CGT CGA ACG GCA GG-3']. The 3+1-63-nt-control DNA fragment was subcloned into the linearized pMD19-RGq-model vector using HiFi DNA Assembly Cloning Kit (New England Biolabs), and then pMD19-model-3+1-63nt control construct was obtained.

pMD19-RGq-model-control-3+1-100nt:
The 3+1-100nt-control insert DNA was prepared by PCR using pUC19 vector as a template with a forward primer [5'-CCT AGA TTA ATG CAA TTC GTG TGA AGA

TCC TTT TTG ATA ATC TCA TGA C-3’] and a reverse primer [5’-TCA ATT GGG TGC TGG AAA TGA ACT CAC GTT AAG GGA TTT TGG TCA TGA G-3’]. As a vector preparation, pMD19-3+1-63nt-control construct was linearized by PCR, using a forward primer [5’-CAT TTC CAG CAC CCA ATT GAA GCT T-3’] and a reverse primer [5’-ACG AAT TGC ATT AAT CTA GGT CGA C-3’]. The 3+1-100-nt-control DNA fragment was subcloned into the linearized pMD19-3+1-63nt-control vector using HiFi DNA Assembly Cloning Kit (New England Biolabs), and then pMD19-model-3+1-100-nt-control constructs were obtained.

pMD19-RGq-model-control-3+1-140nt:

The 3+1-140nt-control insert DNA was prepared by PCR using pUC19 vector as a template with a forward primer [5’-CCT AGA TTA ATG CAA TTC GTG TGA AGA TCC TTT TTG ATA ATC TCA TGA C-3’] and a reverse primer [5’-TCA ATT GGG TGC TGG AAA TGT TTG ATC TTT TCT ACG GGG TCT GAC GCT CAG-3’]. The 3+1-140-nt-control DNA fragment was subcloned into the linearized pMD19-3+1-63nt-control vector using HiFi DNA Assembly Cloning Kit (New England Biolabs), and then pMD19-model-3+1-140-nt-control constructs were obtained.

**Preparation of pIRES-model-5’UTR-FL-RL constructs**

Six model-5’UTR DNA fragments (2+2-63nt, 2+2-100nt, 2+2-140nt, 3+1-63nt, 3+1-100nt, 3+1-140nt) were amplified by PCR using pMD19-RGq-model constructs as templates, with a forward primer [5’-CTA GCC TCG AGA ATT GAT TAG CAT ACG CTA CTG C-3’] and a reverse primer [5’-CCA TGG TGG CGA ATT CTG AAT AGC TTG CAA GTC AT-3’]. Each PCR product was subcloned into the *Eco*R I site in pIRES-FL-RL construct by using HiFi DNA Assembly Cloning Kit (New England Biolabs), and then pIRES-model-5’UTR-2+2-63nt-FL-RL, pIRES-model-5’UTR-100nt-FL-RL, and pIRES-model-5’UTR-140nt-FL-RL constructs and pIRES-model-5’UTR-3+1-63nt-FL-RL, pIRES-model-5’UTR-100nt-FL-RL, and pIRES-model-5’UTR-140nt-FL-RL constructs were obtained.

**Construction of Staple oligomer and siRNA expression vectors**

pAAV-TRPC6-Staple:

The dsDNA fragments of the 26-nt and 40-nt Staple oligomers were prepared by annealing with synthetic oligonucleotides [5’-GAG AAA AGC CTC TAG GCG CAG ACA GGC GGT GGA AGT CAC TAT TTT TTC TAG TGA TAT

CGA TA-3’] and [5’-TAT CGA TAT CAC TAG AAA AAA TAG TGA CTT CCA CCG CCT GTC TGC GCC TAG AGG CTT TTC TC-3’], and [5’- GAG AAA AGC CTC TAG ACC GGA CCG CGC AGA CAG GCG GTG GAA GTC ACT AGT TAG GGT TTT TTC TAG TGA TAT CGA TA -3’] and [5’- TAT CGA TAT CAC TAG AAA AAA CCC TAA CTA GTG ACT TCC ACC GCC TGT CTG CGC GGT CCG GTC TAG AGG CTT TTC TC -3’], respectively. Each dsDNA was subcloned into the *Xba* I-*Spe* I site in a pAAV-U6-ZsGreen1 vector (Takara Bio) using HiFi DNA Assembly Cloning Kit (New England Biolabs), and then pAAV-TRPC6-Staple-26nt and -40-nt constructs were obtained.

pSuper-TPM3-Staple:

The dsDNA fragment of the 26-nt Staple oligomer was prepared by annealing with synthetic oligonucleotides [5’-GAT CCC CCC GGA ACT CAC CAG CTA CTG CTC GCG TTT TTA-3’] and [5’-AGC TTA AAA ACG CGA GCA GTA GCT GGT GAG TTC CGG GGG-3’]. The dsDNA was subcloned into the *Bgl* II-*Hind* III site in pSuper neo vector (Oligoengene) using DNA Ligation Kit (TaKaRa), and then pSuper-TPM3-Staple-26-nt construct was obtained.

pAAV-siRNA:

The dsDNA fragments of the mouse TRPC6 siRNA-1, siRNA-2 and siRNA-3, were prepared by DNA polymerase reaction, using synthetic oligonucleotides: siRNA-1; [5’-GAG AAA AGC CTC TAG GTC ATT CCC TCA ATG TTA ACT GTG AAG CCA CAG ATG GG-3’] and [5’-TAT CGA TAT CAC TAG AAA AAA GTC ATT CCC TCA ATG TTA ACC CAT CTG TGG CTT CAC AG-3’], siRNA-2; [5’-GAG AAA AGC CTC TAG GCA AGG AGC TCA GAA GAT TAC TGT GAA GCC ACA GAT GGG-3’] and [5’-TAT CGA TAT CAC TAG AAA AAA GCA AGG AGC TCA GAA GAT TAC CCA TCT GTG GCT TCA CAG-3’], and siRNA-3; [5’-GAG AAA AGC CTC TAG GCT AGC AGA GCT CAT TAG AAC TGT GAA GCC ACA GAT GGG-3’] and [5’-TAT CGA TAT CAC TAG AAA AAA GCT AGC AGA GCT CAT TAG AAC CCA TCT GTG GCT TCA CAG-3’], respectively. Each dsDNA was subcloned into the *Xba* I-*Spe* I site in pAAV-U6-ZsGreen1 vector (Takara Bio) by using HiFi DNA Assembly Cloning Kit (New England Biolabs), and then pAAV-siRNA-1, -2, and -3 constructs were obtained.

**Preparation of RNA templates for ThT fluorescence assay**

The dsDNA for RNA transcription was prepared from pUC19-TRPC6-5’UTR and pUC19-RGq-model constructs by PCR-amplification with a primer set for TRPC6 5’UTR [5’-TAG AGT ACT TAA TAC GAC TCA CTA TAG GG-3’] and [5’-GGC ACA

GTG CCT GGC CGG-3’], and a primer set for RGq model [5’-TAG AGT ACT TAA TAC GAC TCA CTA TAG GG-3’] and [5’-CTG AAT AGC TTG CAA G-3’], respectively. The dsDNAs were transcribed into single stranded RNAs (ssRNAs) using ScriptMAX® Thermo T7 Transcription Kit (TOYOBO). The ssRNAs were purified with After Tri Reagent RNA Clean Up Kit (Chiyoda Science).

**Preparation of RNA templates for StopAssay**

The dsDNA for RNA transcription was prepared from pUC19-TRPC6-5’UTR or pMD19-RGq-model constructs by PCR-amplification with a primers set for TRPC6 5’UTR [5’-TAG AGT ACT TAA TAC GAC TCA CTA TAG GG-3’] and [5’-CAG GTC GAC TCT AGA GGA TCC GCC AGG GTT TTC CCA GTC ACG AC-3’], and a primer set for RGq model [5’-TAG AGT ACT TAA TAC GAC TCA CTA TAG GG-3’] and [5’-TGG GTA ACG CCA GGG-3’], respectively. The dsDNA was transcribed to ssRNA using ScriptMAX® Thermo T7 Transcription Kit (TOYOBO). The ssRNAs were purified with After Tri Reagent RNA Clean Up Kit (Chiyoda Science).

**Preparation of RNA templates for in vitro translation**

The dsDNA for RNA transcription was prepared from each pIRES construct by PCR-amplification with a primer set for TRPC6 5’UTR [5’-GCT AGA GTA CTT AAT ACG ACT CAC TAT AGG GCT AGC C-3’] and [5’-TTA CTG CTC GTT CTT C-3’], and a primer set for model 5’UTR [5’-TAA TAC GAC TCA CTA TAG GGC  TAG CCT CGA G-3’] and [5’-CTC GAC GCG TGA ATT TTA CAC GGC GAT CTT GCC-3’], respectively. The ssRNAs were transcribed from the dsDNA templates using ScriptMAX® Thermo T7 Transcription Kit (TOYOBO). The ssRNAs were purified with After Tri Reagent RNA Clean Up Kit (Chiyoda Science).

**a**

| Names | RGq control sequences |
|---|---|
| 2+2-63nt | —**AAA**U**AAA**UGUCGACCUAGAUUAAUGCAAUU— 63nt —UUCCAGCACCCAAUUGAAGCUUU**AAA**U**AAA**— |
| 2+2-100nt | —**AAA**U**AAA**UGUCGACCUAGAUUAAUGCAAUU— 100nt—UUCCAGCACCCAAUUGAAGCUUU**AAA**U**AAA**— |
| 2+2-140nt | —**AAA**U**AAA**UGUCGACCUAGAUUAAUGCAAUU— 140nt—UUCCAGCACCCAAUUGAAGCUUU**AAA**U**AAA**— |
| 3+1-63nt | —**AAA**U**AAA**U**AAA**UGUCGACCUAGAUUAAUGCAAUU— 63nt —UUCCAGCACCCAAUUGAAGCUUU**AAA**— |
| 3+1-100nt | —**AAA**U**AAA**U**AAA**UGUCGACCUAGAUUAAUGCAAUU—100nt—UUCCAGCACCCAAUUGAAGCUUU**AAA**— |
| 3+1-140nt | —**AAA**U**AAA**U**AAA**UGUCGACCUAGAUUAAUGCAAUU—140nt—UUCCAGCACCCAAUUGAAGCUUU**AAA**— |

**b**

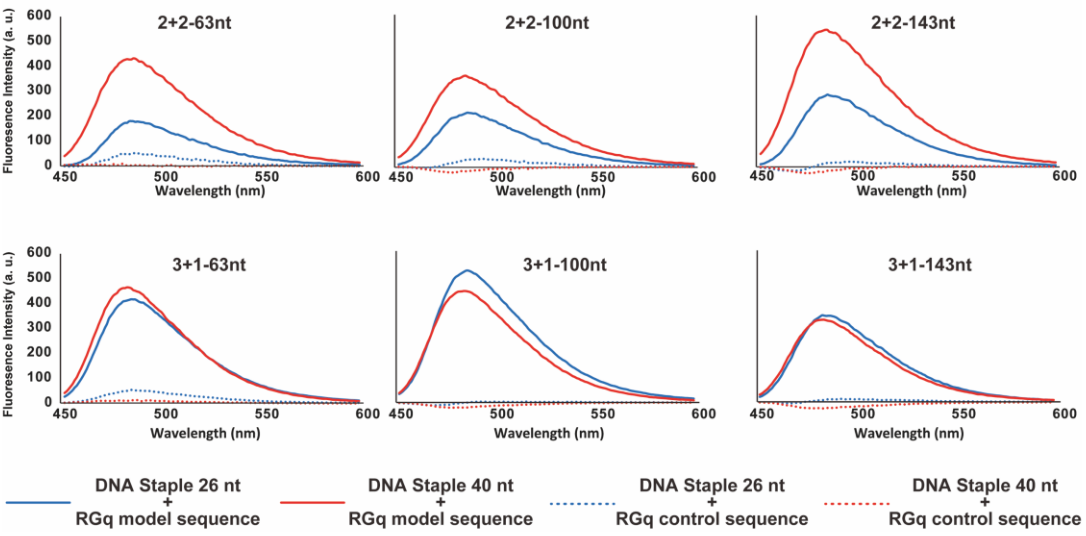


**Figure S1 Evaluation of RGq formation using ThT fluorescent probe**. **a.** The sequences of RGq control. The A-tracts are shown in blue. Staple oligomer recognition sites are underlined. **b.** The blue and red curves show the fluorescence emission spectra of ThT in the presence of 26-nt or 40-nt DNA Staple oligomers, respectively. The solid and dashed curves represent the model sequence with G-tracts and the control with A-tracts, respectively.

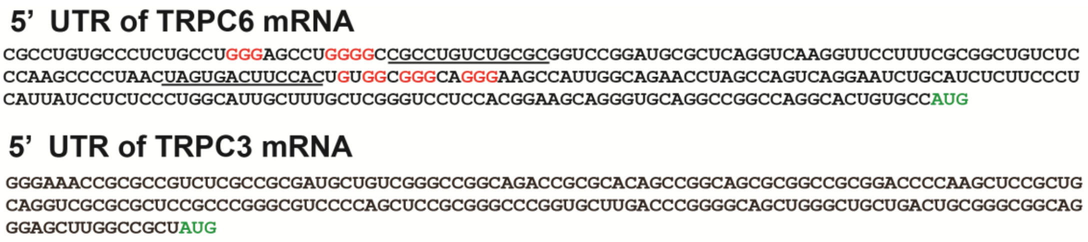


**Figure S2 Nucleotide sequences of 5'UTR of TRPC6 and TRPC3 mRNA.** Guanine repeat sequences and start codon (AUG) are shown in red and green, respectively. Underline shows a hybridization site for the 26-nt RNA Staple oligomer.

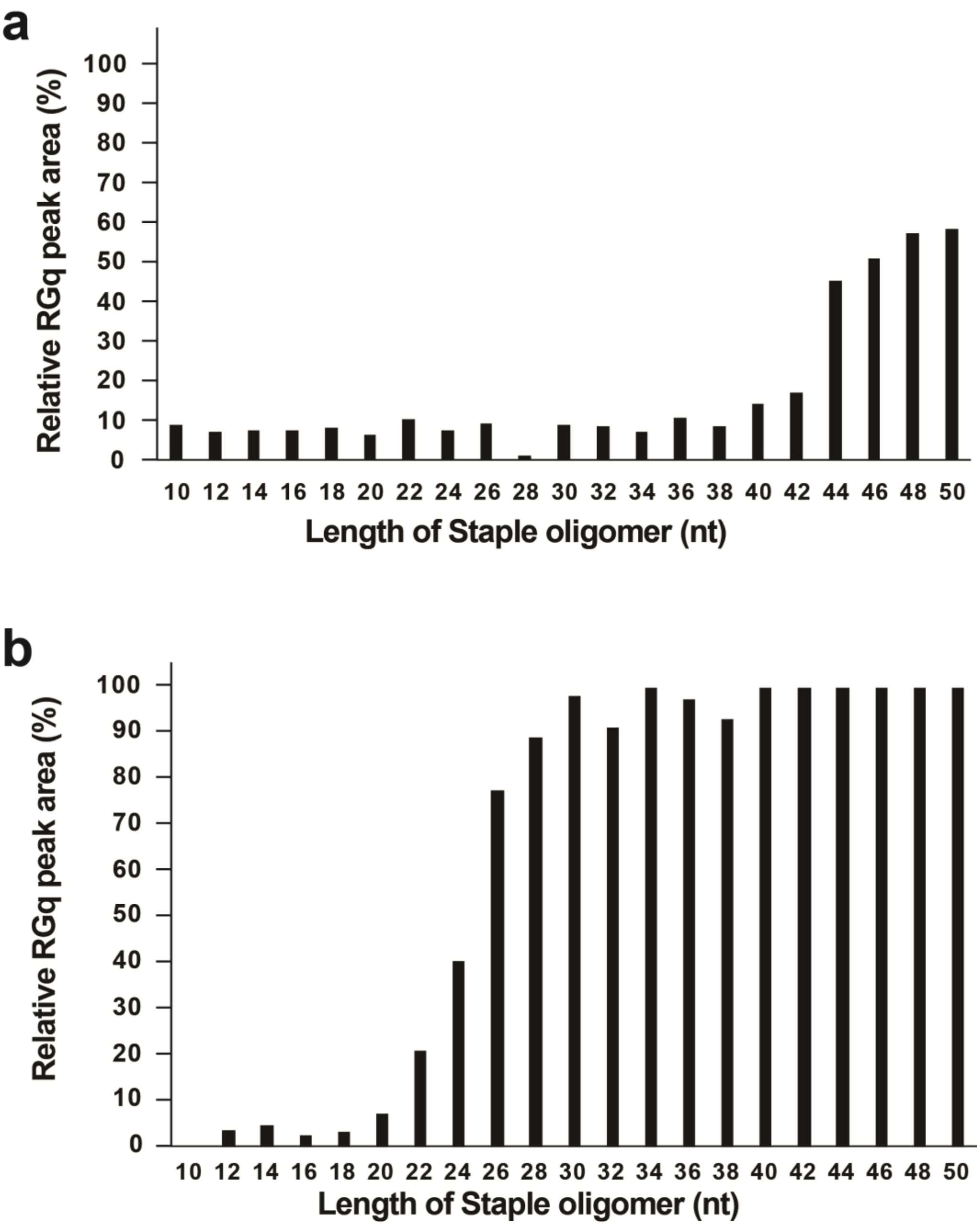


**Figure S3 RGq induction with RNAh technology in various Staple oligomers. a.** Length optimization of DNA Staple. The 44-nt DNA Staple induced approximately 50% RGq structure formation in the target sequence of TRPC6 mRNA. **b.** Length optimization of RNA Staple. The 26-nt RNA Staple induced approximately 80% RGq formation in the target sequence of TRPC6 mRNA. The 44 nt for DNA and the 26 nt for RNA were the minimum length to maintain effective Staple activity in the TPCR6 mRNA.

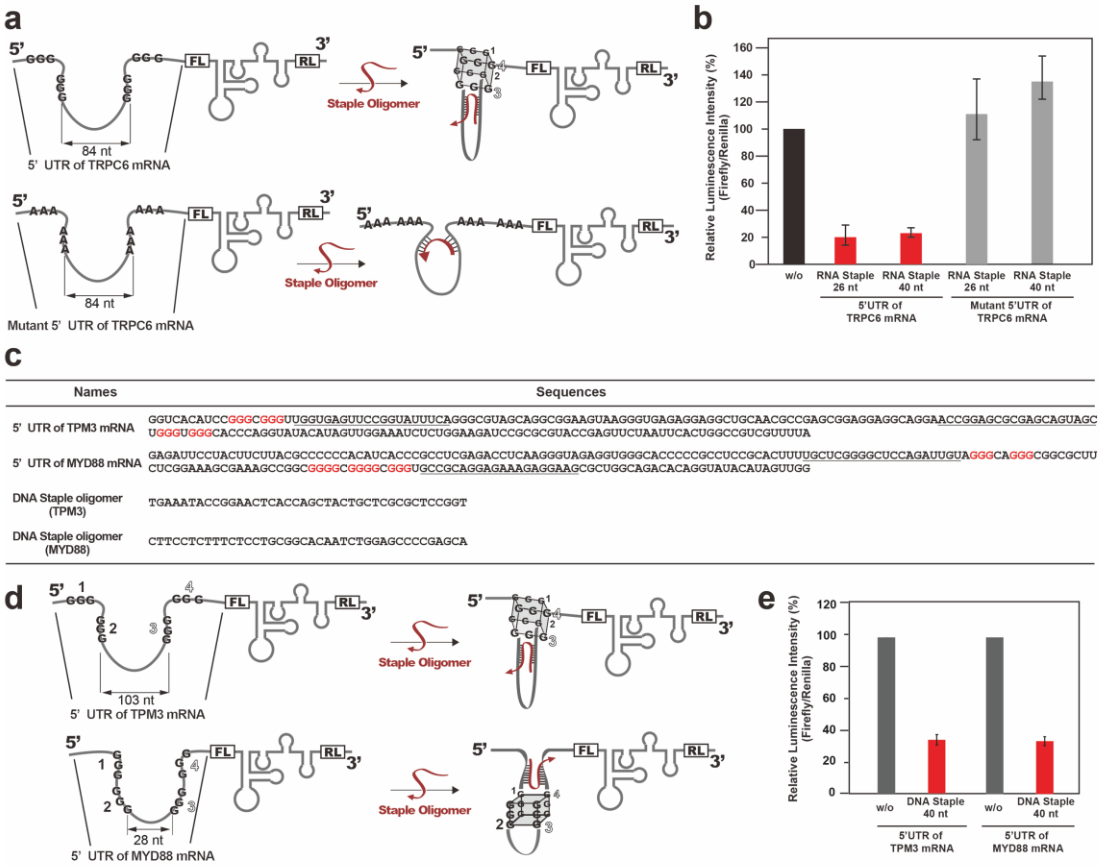


**Figure S4 Validation of the versatility of RNAh technology with Staple oligomers using in vitro translation assay. a**. The mRNAs encoding firefly luciferase (FL) with 5' UTR of TRPC6 (upper scheme) and its mutant (lower scheme) are illustrated. Renilla luciferase (RL) was placed downstream of IRES as an internal expression control. **b**. Translation efficiency was demonstrated as the relative luciferase activity (Firefly/Renilla) for each mRNA with RNA Staple oligomers. Data were normalized by the relative luciferase activity without Staple oligomer. FL expression was strongly repressed by Staple oligomers only in the 5' UTR of TRPC6 but not in its mutant. **c**. Nucleotide sequences of the 5'UTR of TPM3 and MYD88 mRNAs and DNA Staple oligomers. The G-tracts are shown in red. Underline shows a hybridization site for the DNA Staple oligomer. **d**. Illustration of mRNAs encoding FL with 5' UTRs of human cancer-related genes (TPM3; upper scheme and MYD88; lower scheme). **e**. Translation efficiency in each mRNA with the 5'UTR of TPM3 or MYD88, was demonstrated as the relative luciferase activity (Firefly/Renilla) for each mRNA with or without the 40-nt DNA Staple oligomers. Data were normalized by relative luciferase activity without Staple oligomer. The RNA Staple oligomers effectively suppressed in vitro translation of their target mRNAs. Data are mean ± S.D. of three independent experiments.

1

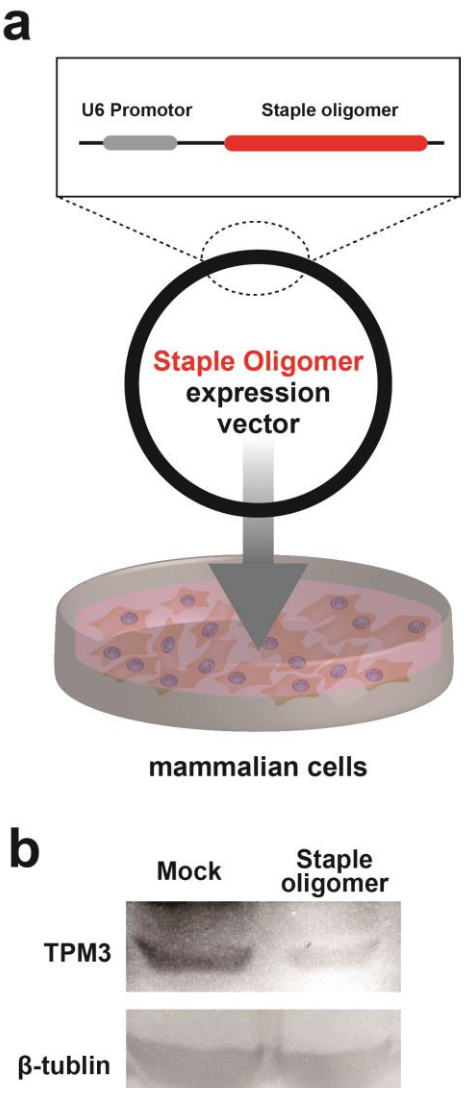


**Figure S5 Effect of RNAh technology on target protein expression in mammalian living cells.** **a.** Design and construction of an expression vector encoding Staple oligomer. The short hairpin RNA expression vector was used to express an RNA Staple oligomer. U6 promotor and Staple oligomer are shown in gray and red, respectively. **b.** Evaluation of the effect of RNA Staple oligomers on TPM3 expression in MCF7 cells by Western blotting. RNA Staple oligomers effectively suppressed TPM3 gene expression.

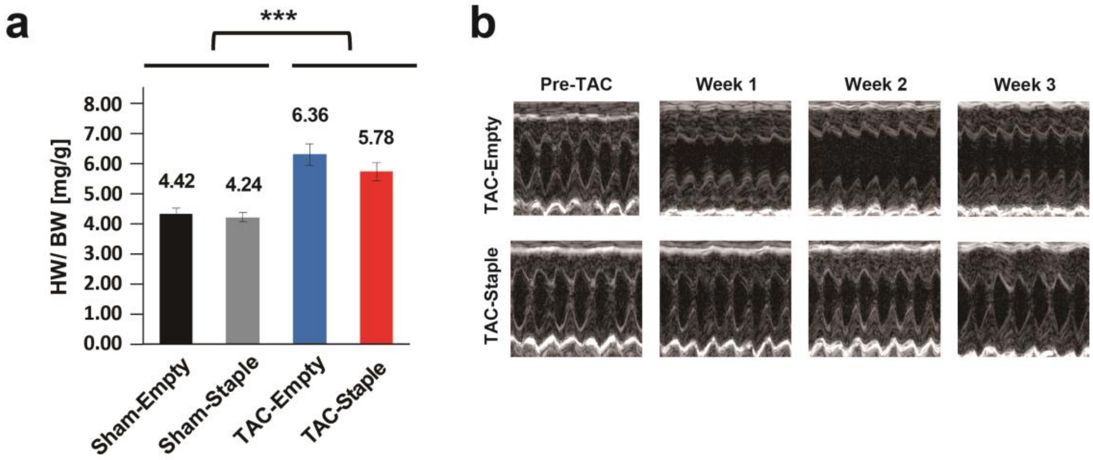


**Figure S6 Effects of Staple oligomers on myocardial hypertrophy in the TAC-treated mouse hearts. a.** The hypertrophic effect of myocardium on heart weight in the TAC-treated and non-TAC-treated groups and in the Staple oligomer-treated and non-Staple oligomer-treated groups. HW/BW represents heart weight/body weight. Whereas the heart weight was increased by TAC treatment (TAC-Empty), the heart weight was significantly suppressed by expression of Staple oligomers (TAC-Staple). Data are shown as mean ±SD. Statistical significance was determined by Student’s t-test: ***$P<0.001$, compared with Sham-treated mice. **b.** Echocardiography showed no significant decrease in cardiac function after TAC treatment in the Staple oligomer-treated mice (lower panel), but a marked decrease in FS values in the non-Staple oligomer-treated mice (upper panel).

a

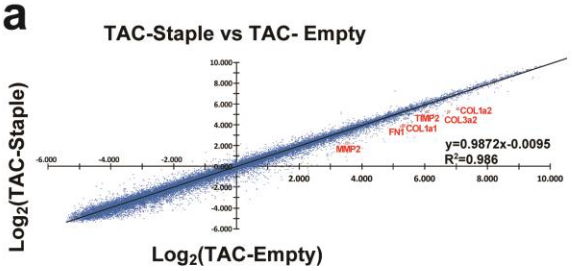


b

| Gene Symbol | Fold Change in microarray (TAC-Staple / TAC-Empty) |
|---|---|
| COL1a1 | 0.48 |
| COL1a2 | 0.52 |
| COL3a1 | 0.33 |
| FN1 | 0.45 |
| MMP2 | 0.42 |
| TIMP2 | 0.55 |

c

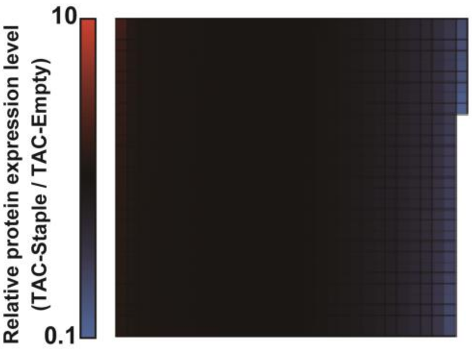


**Figure S7 Microarray and proteomics analysis on myocardial hypertrophy. a**. Scatter plot analysis of microarray from TAC-Empty and TAC-Staple. No detectable change was observed in the expression levels of TRPC6 mRNA with the treatment of Staple oligomers. **b**. The expression levels of cardiac fibrosis-related genes were significantly reduced with Staple oligomer treatments in TAC-treated mice. **c**. Heatmap visualization of protein expression profiles with or without Staple-oligomer treatment in TAC-treated mice.
